# Functional brain networks are associated with both sex and gender in children

**DOI:** 10.1101/2023.11.12.566592

**Authors:** Elvisha Dhamala, Dani S. Bassett, B.T. Thomas Yeo, Avram J. Homes

## Abstract

Sex and gender are associated with human behavior throughout the lifespan and across health and disease, but whether they are associated with similar or distinct neural phenotypes is unknown. Here, we demonstrate that, in children, sex and gender are uniquely reflected in the intrinsic functional connectivity of the brain. Unimodal networks are more strongly associated with sex while heteromodal networks are more strongly associated with gender. These results suggest sex and gender are irreducible to one another not only in society but also in biology.

## Main

Over the last two decades, the interactions between sex, neurobiology, and behavior have been extensively researched^1-9^. However, these studies often report contradictory findings and fail to replicate^7,10^. The growing literature on sex differences^11^ and the lack of reproducibility of many of those reported differences^7^ suggest a potential bias and/or misunderstanding in how we study, interpret, and report findings related to sex. More recently, researchers have begun to question whether these observed differences between males and females are driven by biology (e.g., sex) or if they are a manifestation of social constructs (e.g., gender)^7,10^. In fact, the reality is more complicated in that sex and gender are both influenced by biological and social factors^12,13^. Here, we use the term sex to indicate features of an individual’s physical anatomy, physiology, genetics, and/or hormones at birth, and we use the term gender to indicate features of an individual’s attitude, feelings, and behaviors^14^. Biomedical research thus far has principally focused on understanding the influence of sex on brain and behavior. As such, the contributions of gender are largely unknown.

A fundamental aspect of our human experience is our sex and gender, how we perceive them, and how they are perceived by others. Sex and gender can explain our behavior and influence our health and disease throughout the lifespan. Women, people assigned female at birth (AFAB), and sex/gender minorities have historically been excluded from biomedical research^15,16^. Consequently, this group of individuals is more likely to be underdiagnosed or misdiagnosed for common brain disorders (e.g., ADHD), and suffer adverse effects from treatment interventions (e.g., medications). In the brain sciences, there exist sex and gender differences in the prevalence and expression of psychiatric illnesses and treatment-seeking behaviors. AFAB people are more likely to meet criteria for mood and anxiety disorders, while people assigned male at birth (AMAB) are more likely to be diagnosed with substance use and attention deficit disorders^17^. AFAB people are more likely to report mood problems and seek treatment for mental illnesses^18^. In recent years, researchers have sought to relate these differences in the presentation of psychiatric illnesses to patterns of functional brain organization ^5,19-24^. However, work in this area has largely operated with the assumption that the observed differences are a product of sex, not gender. An understanding of the unique functional brain correlates of sex and gender is essential for the study of brain-related illnesses that exhibit differences across males and females.

Leveraging neuroimaging data from the Adolescent Brain Cognitive Development (ABCD) Study^25^ (4757 children, 2351 females, 9-10 years old) at baseline, and self- and parent-reported gender data at the 1-year follow-up time point, we first evaluated sex differences in gender scores (Figure 1A for all subjects, Supplemental Materials Extended Data Figure 1 for data split by site). Self-reported gender scores measured felt-gender, gender expression, and gender contentedness, while parent-report gender scores measured sex-typed behavior during play and gender dysphoria (Supplemental Materials Extended Data Table 1). Across both self- and parent-report gender measures, higher scores indicate greater sex congruence, which refers to the extent to which an individual’s gender aligns with their assigned sex. Self- and parent-reported gender scores were more similar in children assigned female at birth (AFAB; Spearman correlation, ρ=0.173, p<1.00×10^-10^) than in children assigned male at birth (AMAB; ρ=0.1079, p=8.97×10^-5^), and AMAB children exhibited greater sex congruence than AFAB children for both parent-report (Mann Whitney U statistic, U=1.36×10^6^, p<1.00×10^-10^) and self-report (U=2.13×10^6^, p<1.00×10^-10^). These trends are in line with those previously reported in the entire ABCD sample^14^, indicating that the subsample with neuroimaging data used in these analyses is representative of the full cohort. Extant literature suggests that AMAB children feel more pressure to conform to gender norms than AFAB children^26,27^. This may, in part, explain our results, in which AMAB children report stronger sex-congruent genders than AFAB children.

**Figure 1:**
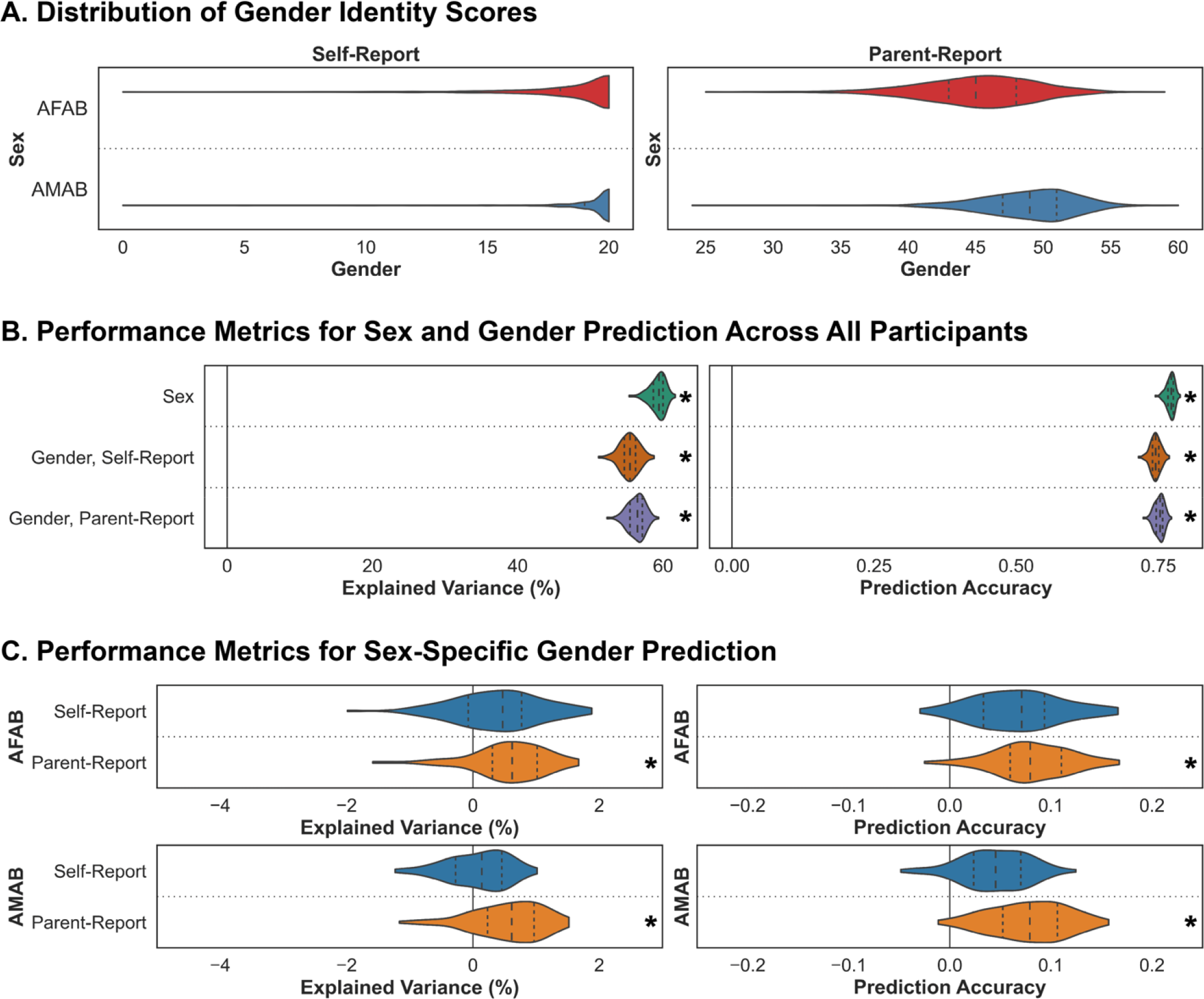
Functional connectivity is associated with assigned sex and gender. (A) Violin plots display the distribution of the self- and parent-reported gender expression scores for AFAB (red) and AMAB (blue) children. (B) Explained variance (%) and prediction accuracy (correlation between true and predicted values) obtained from the models trained to predict sex. Black asterisks (*) indicate that the model performed significantly better than the null models (p<0.05). Results are shown for both the true (green) and the null (orange) predictions. (C) Explained variance (%) and prediction accuracy (correlation between true and predicted values) obtained from the models trained to predict self- and parent-reported gender expression. Black asterisks (*) denote that the model performed significantly better than the null models (corrected p<0.05). For all violin plots, the shape indicates the entire distribution of values; the dashed lines indicate the median; and the dotted lines indicate the interquartile range. AFAB – assigned female at birth; AMAB – assigned male at birth.

Using cross-validated linear ridge regression models, we quantified the associations between functional connectivity and sex as well as gender. Across all individuals, functional connectivity explained 59.27% (p<1.00 ×10^-10^; prediction accuracy, r=0.77, p<1.00 ×10^-10^) of the variance in sex, 55.37% (p<1.00 ×10^-10^; r=0.75, p<1.00 ×10^-10^) of the variance in self-reported gender, and 56.30% (p<1.00 ×10^-10^; r=0.75, p<1.00 ×10^-10^) of the variance in parent-reported gender (Figure 1B). These predictions of gender across all individuals are likely to be confounded by sex (and vice versa), as sex and gender are undeniably related to one another.

To further disentangle the functional correlates of sex from those of gender, we quantified sex-specific associations between functional connectivity and gender. These models were trained separately in AFAB or AMAB children to predict gender. While our models did not successfully predict the self-reported gender scores in either sex (all corrected p-values>0.05), functional connectivity did explain 0.56% (corrected p=0.037; r=0.08, corrected p=0.033) and 0.55% (corrected p=0.037; r=0.08, corrected p=0.033) of the variance in parent-reported gender scores in AFAB and AMAB individuals, respectively (Figure 1C). Detailed model performance metrics and corrected p-values are reported in Table 1. Multiple studies have shown that functional connectivity accurately captures variance related to sex^20,28,29^. Here we replicate those findings in children and further demonstrate that functional connectivity is also associated with parental reports of their children’s gender.

**Table 1:** Detailed performance metrics from models trained to predict sex and gender. Model performance metrics from models trained to predict sex or gender based on functional connectivity. For explained variance (%) and prediction accuracy (correlation between true and predicted measures), mean ± standard deviation performance measured in the hold-out test set across the 100 train/test splits are shown. Model performance metrics were compared to those obtained from null distributions using an exact test for differences to evaluate whether they performed better than chance. Corrected p-values for those comparisons are denoted in parentheses. Models that performed better than chance are indicated by bold type. AFAB – assigned female at birth; AMAB – assigned male at birth.

| Data | Explained Variance (%) |  | Prediction Accuracy |  |
| --- | --- | --- | --- | --- |
|  | AFAB | AMAB | AFAB | AMAB |
| **Assigned Sex** | **59.273 ± 1.077**<br>**(<1.00x10<sup>-10</sup>)** |  | **0.771 ± 0.007**<br>**(<1.00x10<sup>-10</sup>)** |  |
| **Gender Self-Report** | **55.373 ± 1.300**<br>**(<1.00x10<sup>-10</sup>)** |  | **0.745 ± 0.009**<br>**(<1.00x10<sup>-10</sup>)** |  |
| **Gender Parent-Report** | **56.297 ± 1.170**<br>**(<1.00x10<sup>-10</sup>)** |  | **0.752 ± 0.008**<br>**(<1.00x10<sup>-10</sup>)** |  |
| **Gender Self-Report** | 0.393 ± 0.693<br>(0.162) | 0.077 ± 0.483<br>(0.198) | 0.068 ± 0.044<br>(0.136) | 0.044 ± 0.033<br>(0.147) |
| **Gender Parent-Report** | **0.563 ± 0.596**<br>**(0.037)** | **0.551 ± 0.557**<br>**(0.037)** | **0.081 ± 0.038**<br>**(0.033)** | **0.078 ± 0.037**<br>**(0.033)** |

The Haufe transformation^30^ was applied to the feature weights extracted from the models to increase their interpretability and reliability^31^, and the absolute Haufe-transformed weights were averaged to compute a mean absolute feature importance score (at the regional pairwise level). We evaluated the correlations between the features extracted from the different prediction models (Figure 2A). For the sex-independent gender prediction models, functional connections associated with sex largely overlapped with those associated with gender (r_self-report_=1.00, r_parent-report_=0.99), suggesting that models trained to capture variability in gender are capturing variance related to sex, and vice versa. For the sex-specific models, functional connections associated with sex were distinct from the functional connections associated with gender in AFAB (r_self-report_=0.15, r_parent-report_=0.13) and AMAB (r_self-report_=0.12, r_parent-report_=0.11) children. Functional connections associated with gender were weakly correlated across the sexes for the self-report measures (r=0.30) but distinct for the parent-report measures (r=0.18). Finally, functional connections associated with gender were moderately correlated across the self- and parent-report measures in AFAB children (r=0.46) but uncorrelated in AMAB children (r=0.19). These results suggest that sex and gender, although strongly correlated, are uniquely represented in functional networks.

**Figure 2:**
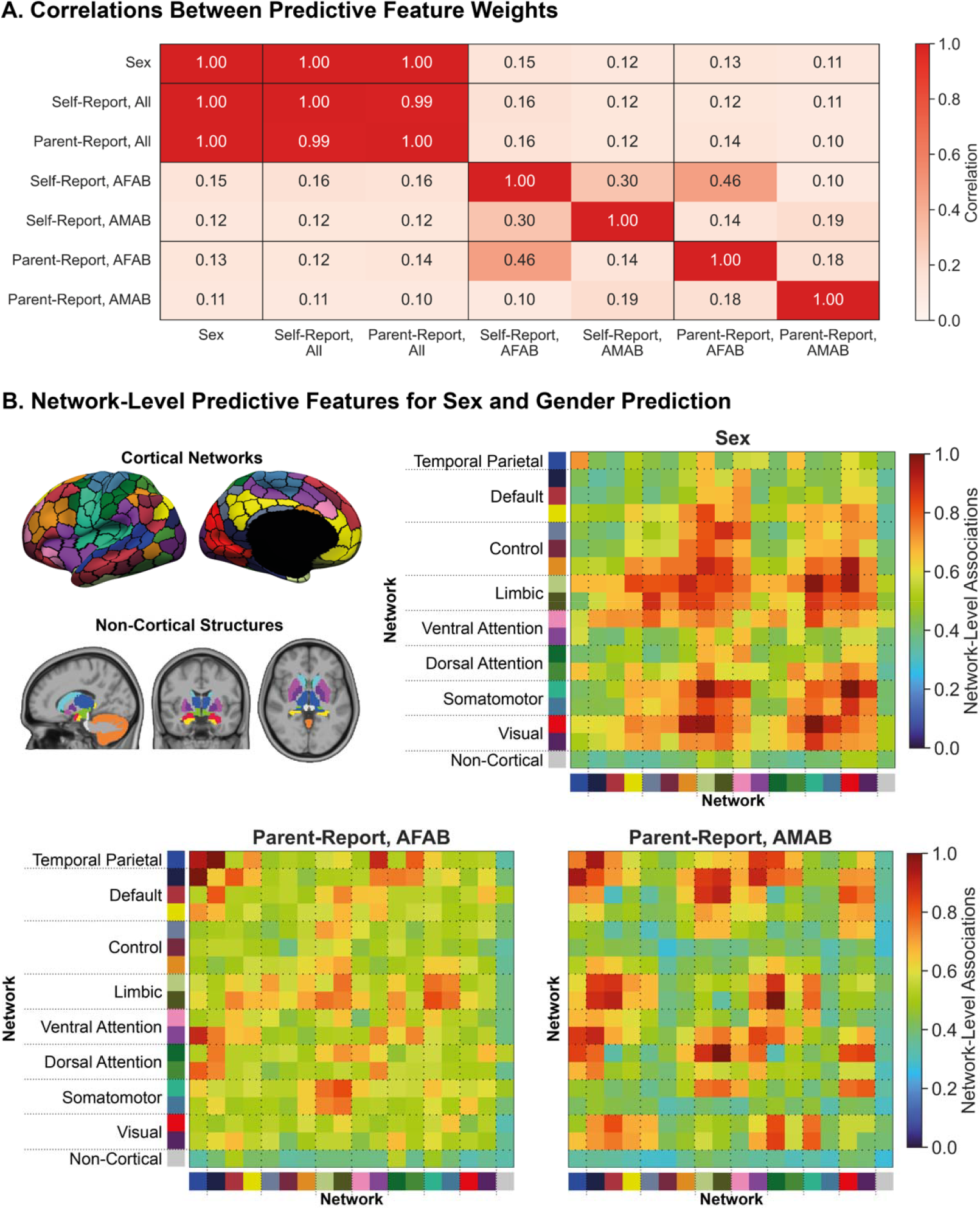
Distinct functional networks are associated with assigned sex and gender. (A) Correlation coefficient between Haufe-transformed absolute pairwise regional feature weights from distinct models trained to predict assigned sex and gender expression. Models trained to predict gender were either trained across all participants (All), only in AFAB children (AFAB), or only in AMAB children (AMAB). Warmer colors indicate a stronger correlation between the feature weights. (B) Regional pairwise feature weights were summarized to a network-level by mapping the Schaefer 400 cortical parcels to 17 large-scale cortical networks and assigning the non-cortical regions to a single non-cortical network (top left). Cortical network image reproduced with permission from https://doi.org/10.6084/m9.figshare.10062482.v1 and non-cortical network image reproduced with permission from https://doi.org/10.6084/m9.figshare.10063016.v1 under a CC BY 4.0 license. Associations between functional network connectivity and sex (top right) and parent-reported gender expression (bottom) are shown as per the colormap, where warmer colors indicate stronger correlations and cooler colors indicate weaker correlations. AFAB – assigned female at birth; AMAB – assigned male at birth.

Finally, we established the network-level functional correlates of sex and gender by mapping regional feature weights onto 17 large-scale cortical networks^32^ and one non-cortical network (Figure 2B). Here, we focus our analyses on the overlap between the functional correlates of sex and of sex-specific parent-reported measures of gender, as the sex-independent correlates of self- and parent-reported measures of gender (Supplemental Materials Extended Figure 2) were nearly identical to those of sex, and the functional correlates of sex-specific self-reported measures (Supplemental Materials Extended Figure 3) correspond to models that did not perform better than chance. Functional correlates of sex are largely found in the somatomotor, visual, control and limbic networks, while the functional correlates of gender are more dispersed throughout the cortical networks. In AFAB children, functional correlates of gender largely involved connections within and between temporal parietal, limbic, dorsal/ventral attention, and somatomotor networks. In AMAB children, functional correlates of gender included connections within and between temporal parietal, default mode, limbic, dorsal/ventral attention, somatomotor, and visual networks. Based on these findings, we can speculate that distinct functional connections within/between unimodal and heteromodal networks are associated with sex and gender.

Sex and gender differences in biology and behavior are tied to health outcomes throughout the lifespan^33^. An understanding of the neurobiological underpinnings of sex and gender is crucial for the subsequent identification of how sex and gender influence health and illness and the development of sex-specific and gender-oriented diagnostic and prognostic tools^13,34^. Here, we demonstrate that functional connectivity is associated with sex and parent-reported gender. Our predictions of gender (beyond sex) are far less accurate than predictions of sex or gender alone, suggesting that gender may be a more complex construct that is not as clearly represented in functional connectivity patterns. Moreover, our inability to capture variance in self-reported gender may be due to the general lack of population-level variability in the self-reported scores relative to the parent-reported scores.

Functional brain networks mature nonuniformly across cortical networks^35-37^ and sexes^2^ during adolescence. Unimodal sensory networks (e.g., visual, auditory, somatosensory) responsible for responding to stimuli within one sensory modality mature first, followed by heteromodal association networks (e.g., dorsal attention, ventral attention, control) involved in higher-order cognitive and social processes. While both unimodal and heteromodal networks are involved in sex and gender predictions, sex is more strongly associated with unimodal networks while gender is more strongly associated with heteromodal networks. Although speculative, the stronger associations between sex and unimodal networks may be due to the delayed development of heteromodal networks in children^38^. Moreover, a distinct set of functional connections are associated with gender (after accounting for sex). In AFAB children, the strongest associations were observed in the temporal parietal and attention networks, whereas in AMAB children, we see a more dispersed pattern of association involving heteromodal networks as well as visual and somatomotor networks. Taken together, these findings suggest the functional correlates of sex are distinct from the functional correlates of gender, and the unique multidimensional constructs that comprise gender are differentially associated with functional connectivity patterns in AFAB and AMAB children. As such, sex and gender must both be studied concurrently to fully capture the differences and similarities that exist between males and females, between boys and girls, and between other genders.

These findings are subject to several limitations. First, sex is not binary. However, in the ABCD sample analyzed here, all subjects reported their sex as either *female* or *male*. As such, we only considered the neural correlates of binary sex and the correlates of gender in AFAB and AMAB children. Additional analyses in a more sex- and gender-diverse sample may reveal further insights. Second, these analyses were performed in a relatively young cohort. As these children undergo puberty and adolescence, their understanding and expression of their gender will likely change. This will be paralleled by changes in functional network organization^36,37^, especially in the heteromodal networks observed to be associated with gender. As such, future analyses should seek to evaluate how these functional network representations of sex and gender change during the transition from childhood to adolescence and across adulthood. Third, gender was assessed at a single time point, and using a set of questions that assumed a static form of gender identity and expression. Future work across multiple time points could instead use questions that allow for the assessment of gender fluidity over a range of time scales. Finally, gender is influenced by local cultural norms and shared societal experiences. The ABCD dataset was collected entirely in the United States and is not representative of the global population^39^. Subsequent analyses should investigate whether similar relationships exist in other countries. For a more detailed discussion of our current findings and our interpretations of them, we refer readers to our Supplemental Discussion.

A comprehensive understanding of the neurobiological correlates of sex and gender is necessary if we are to understand health and disease in sex- and gender-diverse samples. Here, we identify the distinct functional network correlates of assigned sex and gender expression in the developing human brain. How these correlates may be maintained or altered during development and adulthood, and how they may relate to genderfluid experiences at any age, remains to be established.

## Methods

### Dataset

The Adolescent Brain Cognitive Development (ABCD) dataset is a large community-based sample of children and adolescents who were assessed on a comprehensive set of neuroimaging, behavioral, developmental, and psychiatric batteries^25^. In this study, we used minimally preprocessed neuroimaging data acquired at the baseline time point along with self- and parent-reported gender data at the 1-year follow-up time point from the NIMH Data Archive for ABCD Release 2.0.1. Magnetic resonance (MR) images were acquired across 21 sites in the United States of America using harmonized protocols for GE and Siemens scanners. In line with our prior work^40,41^, exclusion criteria were used to ensure quality control. As recommended by the ABCD consortium, we excluded individuals who were scanned using Philips scanners due to incorrect preprocessing (https://github.com/ABCD-STUDY/fMRI-cleanup). For the T1 data, individuals who did not pass recon-all quality control^42^ were removed. For the functional connectivity data, functional runs with boundary-based registration (BBR) costs greater than 0.6 were excluded. Further, volumes with framewise displacement (FD)>0.3 mm or voxel-wise differentiated signal variance (DVARS)>50, along with one volume before and two volumes after, were marked as outliers and subsequently censored. Uncensored segments of data containing fewer than five contiguous volumes were also censored^43,44^. Functional runs with over half of their volumes censored and/or max FD>5mm were removed. Individuals who did not have at least 4 minutes of data were also excluded. Individuals who did not have all self- and parent-reported gender data were also excluded. Next, data from sites with fewer than 50 individuals were also excluded. Finally, we excluded siblings to prevent unintended biases due to inherent heritability in neurobiological and/or behavioral measures. Our final sample comprised 4757 children (2315 assigned female at birth, ages 9-10 years) from the Adolescent Brain Cognitive Development (ABCD) 2.0.1 release^25^.

### Sex and Gender Data

We included sex assigned at birth (referred to as ‘sex’) and gender data from the Youth Self-Report and Parent-Report Gender Questionnaires^14^. All participants included in these analyses completed the Youth Gender Survey which includes 4 questions that measure felt-gender, gender expression, and gender contentedness. Additionally, their parents/caregivers completed an adapted Gender Identity Questionnaire^45,46^ which included 12 questions that measure sex-typed behavior during play and gender dysphoria. A list of all questions asked in the self-report and parent-report surveys can be found in Extended Data Table 1 (Supplemental Materials). We computed summary self-report and parent-report gender scores for all participants by computing the sum across all questions within each questionnaire, respectively, and used these summary scores in our analyses. We used non-parametric Mann-Whitney U rank tests to evaluate sex differences in the gender scores. All p-values were corrected for multiple comparisons using the Benjamini-Hochberg False Discovery Rate (q=0.05) procedure^47^. We also computed non-parametric correlations between the gender scores for each assigned sex to evaluate any underlying relationships that may exist.

### Image Acquisition and Processing

The minimally preprocessed MRI data were processed as previously described^41,48^. Briefly, minimally preprocessed T1 data were further processed using FreeSurfer 5.3.0^49-52^ to generate cortical surface meshes for each individual, which were then registered to a common spherical coordinate system^51,52^. Minimally preprocessed fMRI data were further processed with the following steps: (1) removal of initial frames, with the number of frames removed depending on the type of scanner^42^ and (2) alignment with the T1 images using boundary-based registration^53^ with FsFast. Framewise displacement (FD)^54^ and voxel-wise differentiated signal variance (DVARS)^55^ were computed using fsl_motion_outliers. Respiratory pseudomotion was filtered out using a bandstop filter (0.31-0.43 Hz) before computing FD^56-58^. A total of 18 nuisance covariates were also regressed out of the fMRI time series: global signal, six motion correction parameters, averaged ventricular signal, averaged white matter signal, and their temporal derivatives. Regression coefficients were estimated from the non-censored volumes. Global signal regression was performed as we are interested in behavioral prediction, and global signal regression has been shown to improve behavioral prediction performance^59,60^. Finally, the brain scans were interpolated across censored frames using least squares spectral estimation^61^, band-pass filtered (0.009 Hz ≤ f ≤ 0.08 Hz), projected onto FreeSurfer fsaverage6 surface space, and smoothed using a 6 mm full-width half maximum kernel. Once processed, we extracted regional time series for 400 cortical^62^ and 19 non-cortical^63^ parcels. Full correlations were computed between those time series yielding a 419×419 pairwise regional functional connectivity matrix. All processing as described was completed on a local server.

### Predictive Modeling

Linear ridge regression models avoid overfitting, are interpretable, and are relatively computationally inexpensive compared to deep learning algorithms for brain-based behavioral predictions^64,65^. Here, using a similar framework as those previously used by our research team^22,23,66,67^, we perform novel analyses to establish the functional brain correlates of assigned sex and gender. We used cross-validated ridge regression models to predict sex and gender based on functional connectivity. To facilitate comparisons of model performance and feature contributions across sex and gender predictions, we implemented the linear ridge regression framework for sex predictions instead of a classification model. Models predicting sex included all individuals (AMAB and AFAB), while models predicting gender either included all individuals or were sex-specific (i.e., trained and tested separately for each sex). For models predicting gender that included all individuals, gender scores were reversed in AFAB children such that they ranged from 0 to 20 for self-report scores (with 0 indicating more feminine gender identities and expressions and 20 indicating more masculine gender identities and expressions) and 0 to 60 for parent-report scores (with 0 indicating more feminine gender identities and expressions and 60 indicating more masculine gender identities and expressions). For each model, we split the data into 100 distinct train and test sets (at approximately a 4:1 ratio) without replacement. Imaging site was considered when splitting the data such that we placed all participants from a given site either in the train or test set but not split across the two. Within each train set, we optimized the regularization parameter using three-fold cross-validation while similarly accounting for imaging site as in the initial train-test split. Once optimized, we evaluated models on the corresponding test set. We repeated this process for each of 100 distinct train-test splits to obtain a distribution of prediction accuracy and explained variance. To evaluate model significance, for each set of predictive models, a corresponding set of null models was generated as follows: the output variable was randomly permuted 1000 times, and each permutation was used to train and test a null model using a randomly selected regularization parameter from the set of selected parameters from the original model. Prediction accuracy from each of the null models was then compared to the average accuracy from the corresponding distribution of model accuracies of the original (true) models. The p-value for each model’s significance is defined as the proportion of null models with prediction accuracies or explained variances greater than or equal to those corresponding to the original (true) distributions. All p-values were corrected for multiple comparisons across the gender measures using the Benjamini-Hochberg False Discovery Rate (q=0.05) procedure^47^.

### Feature Weights

We used the Haufe transformation^30^ to transform feature weights obtained from the linear ridge regression models to increase their interpretability and reliability^41,48,68^. For each train split, we used feature weights obtained from the model, W, the covariance of the input data (functional connectivity), Σ_x_, and the covariance of the output data (behavioral score), Σ_y_, to compute the Haufe-transformed feature weights, A, as follows: 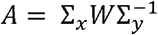. We then averaged the absolute Haufe-transformed feature weights across the 100 splits to obtain a mean feature importance value. We computed full correlations between mean feature importance obtained from the different models to evaluate whether they relied on shared or unique features to predict sex/gender. For all models, we also summarized pairwise regional feature importance at a network-level to support interpretability as previously described^22^. Briefly, cortical parcels were assigned to one of 17 networks from the Yeo 17-network parcellation^32^, and subcortical, brainstem, and cerebellar parcels were assigned to a single non-cortical network for convenience. Regional pairwise absolute feature weights were averaged to yield network-level estimates of associations between functional connectivity and sex/gender.

### Data and Code Availability

All ABCD data can be accessed via the NIMH Data Archive. All code used to generate the results can be found on GitHub (https://github.com/elvisha/sg_predictions).

## Supporting information

Supplemental Discussion

## Acknowledgements

This work was supported by the following awards to ED: the Northwell Health Advancing Women in Science and Medicine Career Development Award and the Feinstein Institutes for Medical Research Emerging Scientist Award. Additional support was provided by the National Institute of Mental Health (R01MH123245 to AJH and R01MH120080 to AJH and BTTY). Additional support was also provided by the following awards to BTTY: NUS Yong Loo Lin School of Medicine (NUHSRO/2020/124/TMR/LOA), the Singapore National Medical Research Council (NMRC) LCG (OFLCG19May-0035), NMRC CTG-IIT (CTGIIT23jan-0001), NMRC STaR (STaR20nov-0003), Singapore Ministry of Health (MOH) Centre Grant (CG21APR1009), the Temasek Foundation (TF2223-IMH-01), and the United States National Institutes of Health (R01MH133334). Any opinions, findings, and conclusions or recommendations expressed here are those of the authors and do not necessarily reflect the views of the funders.

