## Supplemental Materials for "Functional brain networks are associated with both sex and gender in children"

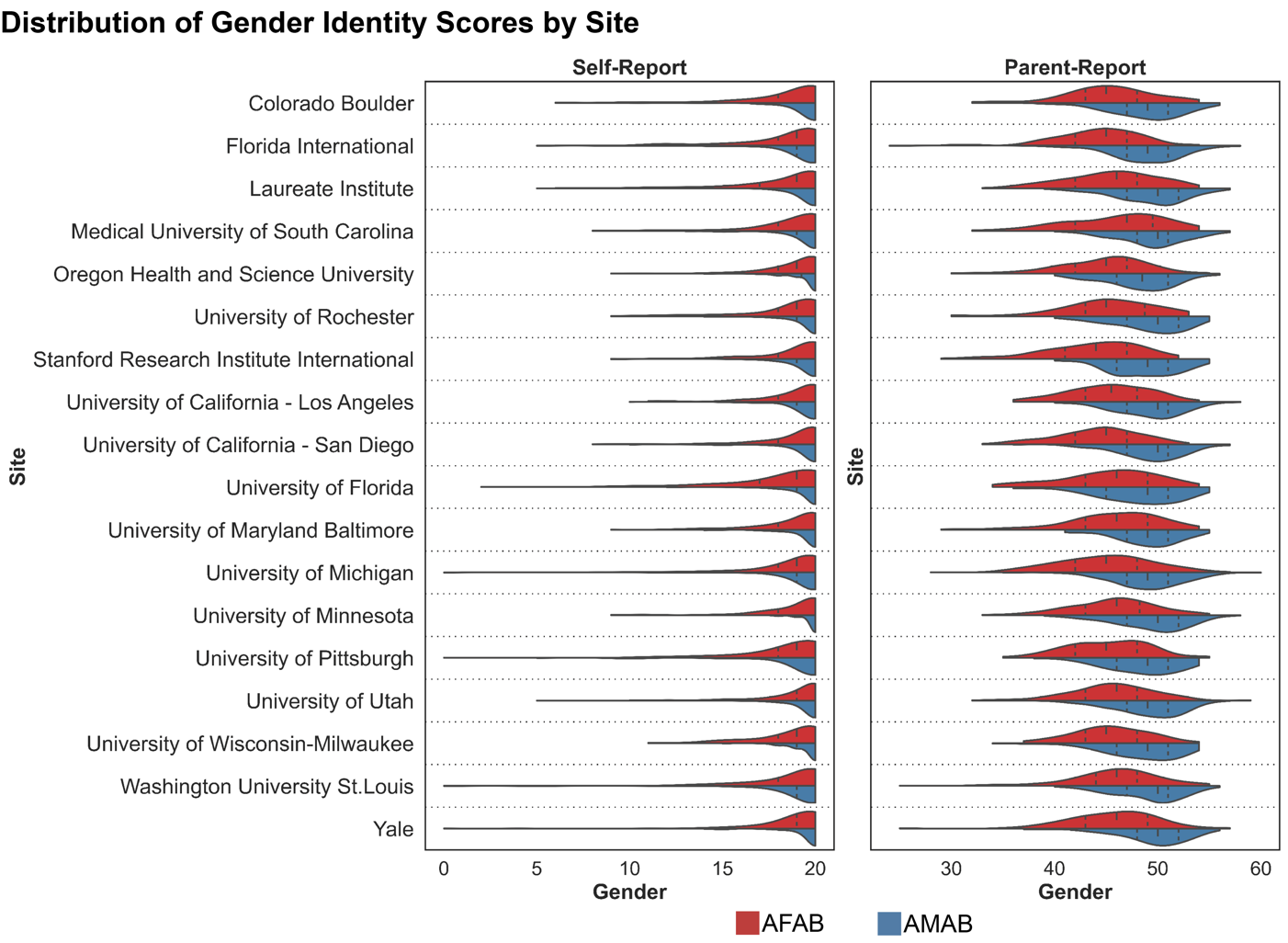
**Extended Data Figure 1: Distribution of Gender Scores by Site**

Violin plots display the distribution of the self- and parent- reported gender scores for AFAB children (red) and AMAB children (blue) for each site. The shape of the violin indicates the entire distribution of values, dashed lines indicate the median, and dotted lines indicate the interquartile range. AFAB – assigned female at birth; AMAB – assigned male at birth.

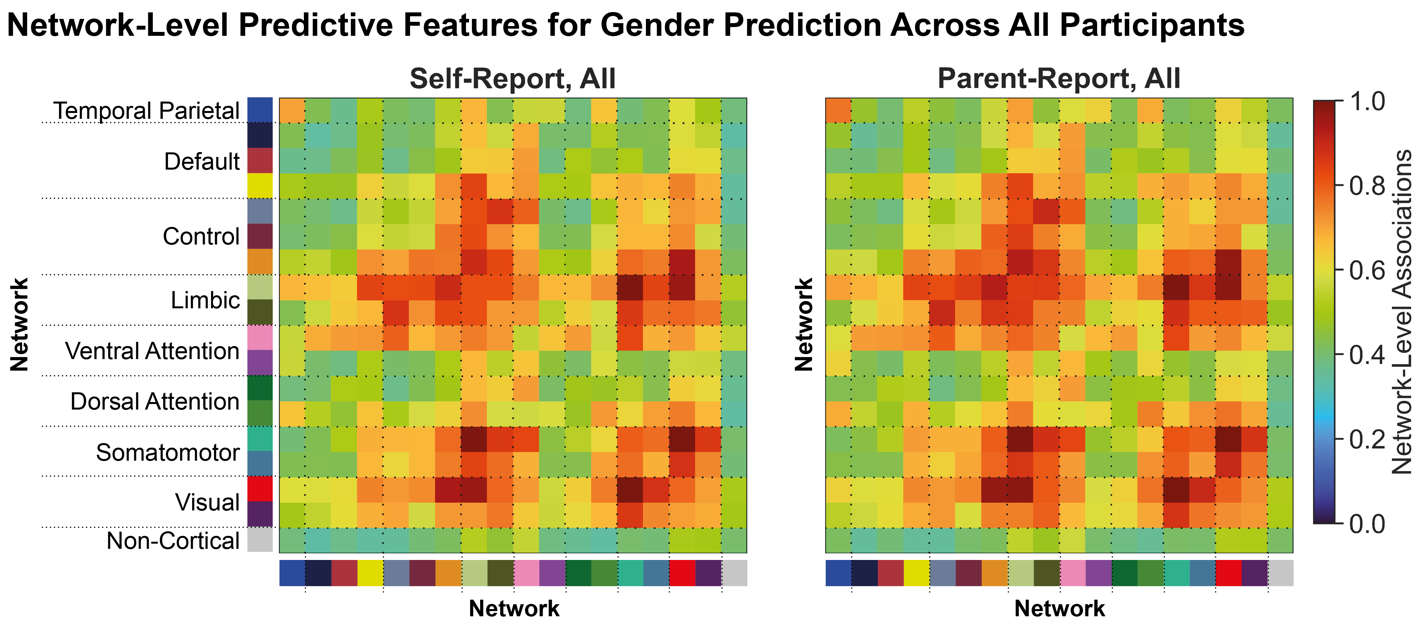

**Extended Data Figure 2: Network-Level Predictive Features for Gender Prediction Across All Participants**

Regional pairwise feature weights to predict gender across all participants were summarized to a network level by mapping the Schaefer 400 cortical parcels to 17 large-scale cortical networks and assigning the non-cortical regions to a single non-cortical network (Figure 2B). Associations between functional network connectivity and self-reported (left) and parent-reported (right) gender are shown as per the colormap, where warmer colors indicate stronger correlations and cooler colors indicate weaker correlations.

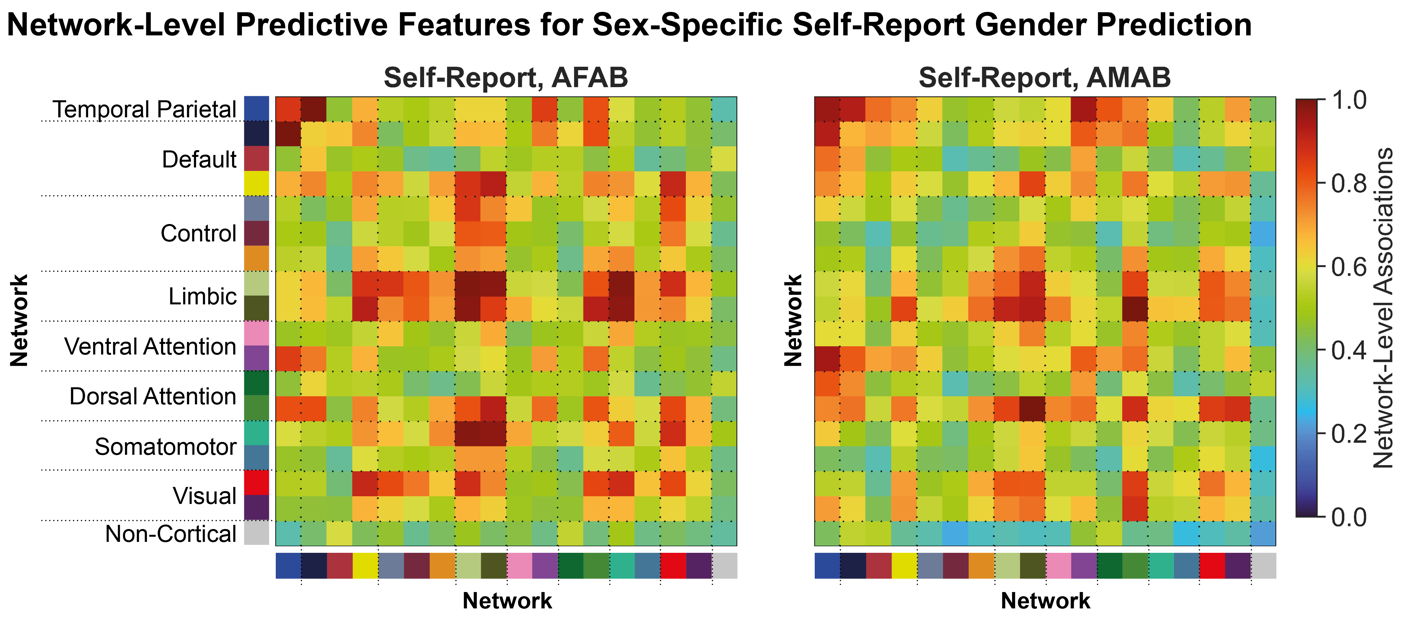

**Extended Data Figure 3: Network-Level Predictive Features for Sex-Specific Self-Report Gender Prediction**

Regional pairwise feature weights to predict self-reported gender in a sex-specific manner were summarized to a network level by mapping the Schaefer 400 cortical parcels to 17 large-scale cortical networks and assigning the non-cortical regions to a single non-cortical network (Figure 2B). Associations between functional network connectivity and self-reported gender in AFAB children (left) and AMAB children (right) are shown as per the colormap, where warmer colors indicate stronger correlations and cooler colors indicate weaker correlations. AFAB – assigned female at birth; AMAB – assigned male at birth.

**Extended Data Table 1: ABCD Gender Survey Questions**

The ABCD Youth (Self-Report) Gender Survey included 4 questions that measured felt-gender, gender expression, and gender contentedness. The ABCD Parent-Report Gender Survey included 12 questions that measured sex-typed behavior during play and gender dysphoria in assigned male at birth (AMAB) and assigned female at birth (AFAB) children.

| **Survey** | **Domain** | **AMAB** | **AFAB** |
| --- | --- | --- | --- |
| Self-Report | Sex-Congruent Felt-Gender | How much do you feel like a boy? | How much do you feel like a girl? |
|  | Sex-Incongruent Felt Gender | How much do you feel like a girl? | How much do you feel like a boy? |
|  | Contentedness | How much have you had the wish to be a girl? | How much have you had the wish to be a boy? |
|  | Expression | How much have you dressed or acted as a girl during play? | How much have you dressed or acted as a girl during play? |
| Parent-Report | Sex-typed behavior during play | His favorite playmates are: | Her favorite playmates are: |
|  |  | He plays with girl-type dolls, such as "Barbie". | She plays with girl-type dolls, such as "Barbie". |
|  |  | He plays with boy-type dolls such as action figures or "GI-Joe". | She plays with boy-type dolls such as action figures or "GI-Joe". |
|  |  | He experiments with cosmetics (makeup) and jewelry. | She experiments with cosmetics (makeup) and jewelry. |
|  |  | He imitates female characters seen on TV or in the movies. | She imitates female characters seen on TV or in the movies. |
|  |  | He imitates male characters seen on TV or in the movies. | She imitates male characters seen on TV or in the movies. |
|  |  | He plays sports with boys (but not girls). | She plays sports with boys (but not girls). |
|  |  | He plays sports with girls (but not boys). | She plays sports with girls (but not boys). |
|  |  | *In playing "mother/father", "house", or "school" games, he takes the role of:* | *In playing "mother/father", "house", or "school" games, she takes the role of:* |
|  |  | He plays "girl-type" games (as compared to "boy-type" games). | She plays "girl-type" games (as compared to "boy-type" games). |
|  |  | *In dress-up games, he likes to dress up as:* | *In dress-up games, she likes to dress up as:* |
|  | Gender dysphoria | He states the wish to be a girl or woman. | She states the wish to be a boy or man. |
|  |  | He states that he is a girl or woman. | She states that she is a boy or man. |
|  |  | He talks about not liking his sexual anatomy (private parts). | She talks about not liking her sexual anatomy (private parts). |
