## Supplemental Discussion for "Functional brain networks are associated with both sex and gender in children"

Scientific studies of sex and gender have historically been intertwined with social and political processes that have consistently harmed marginalized identities. In writing a manuscript in this space, we acknowledge that history, and seek to frame our findings carefully so as to minimize the potential for our work to be used inappropriately or to cause more harm to sex and gender minorities. This supplemental discussion section comprises our attempt at such a framing. In writing it, we have sought to use language that is currently accepted as inclusive. However, we recognize that language changes over time; we hope that future readers will view our discussion in the light of current knowledge and thought, and expand and embrace the most inclusive language of their time in framing and discussing our results.

**Sex**. In the main text, we use the term “sex” to indicate the sex assigned to the child at birth. In contemporary medicine in the United States of America, sex is assigned based on visual inspection of the genitals at birth. In 99.95% of births, sex is assigned as either “male” or “female”; in the other 0.05% of births, sex is assigned as intersex^1^. Historically, these latter cases that did not fit neatly into either “male” or “female” problematized the definition of sex. Medical practitioners and science scholars came to expand the definition of sex to include a variety of additional variables including chromosomes, gonads, hormones, internal genital morphology, rearing, gender role, and sexual orientation^2^. The process of definition, across a variety of eras from the renaissance to today, served to politicize the body in ways that supported flawed societal frames of normality^3^. Moreover, such definitions have and continue to be used to justify medical violence^4^.

**Gender**. Gender has similarly been conceptualized in increasingly complex ways. Gender is a multifaceted, multivalent, and dynamic aspect of an individual’s identity that encompasses their interactions, expressions, behaviors, and socially constructed roles^5^. In the main text, we use the term “gender” to indicate participants’ self- and parent- reported genders as measured by the Youth Gender Survey and an adapted Gender Identity Questionnaire, respectively. Gender is not binary, and we do not treat it as such in our analyses. Throughout history, transgender and non-binary gender identities have been reported across many cultures^6^. As an example, many Indigenous cultures have long recognized the existence of multiple gender identities^7^. In modern history, gendered terms have been wielded as a tool of oppression, reinforcing patriarchal power structures and societal hierarchies, subsequently limiting opportunities and rights for women and gender minorities. In medicine, massive disparities in access to healthcare treatments, and resources, exist due to discrimination based on gender^8^.

**Sex and Gender Can Relate in Complex Ways.** Studies examining the neuroscience of sex/gender have historically sought to identify basic biological differences between (binary) sexes. Sex and gender are often conflated in biomedical research based on the incorrect assumption that they are determined by the same factors and the two are directly related to one another^5^. However, sex and gender are complex multidimensional constructs associated with a host of biological, social, and environmental factors. A recent framework that has been proposed (see Figure 1 in Chapter 1 of ^9^) to understand the neuroscience of gender considers intrinsic and extrinsic influences on gender, which include biological, physiological, psychosocial, and sociocultural factors^9^. Due to the complex nature of these factors and the nonlinear interactions between them, sex and gender should be clearly and carefully defined and distinguished in research. Importantly, we should not reduce one to the other, but understand that both exist in a bidirectional network that helps make an individual who they are.

**Essentialism and Biological Explanations of Gender.** Essentialism is a philosophical concept that suggests that objects, ideas, and even people have inherent and unchanging characteristics that define their true nature. In the context of human identity, essentialism proposes that individuals have innate and immutable qualities that determine their behaviors, abilities, and roles in society^10^. This idea has been applied to gender, in what is referred to as gender essentialism. Gender essentialism posits that there are distinct and innate differences between men and women, and these differences dictate their respective gender roles in society^11^. This notion has often been used to reinforce traditional gender norms and stereotypes in society, perpetuating gender inequality and limiting opportunities^12^. Essentialism and gender essentialism oversimplify the complex nature of human identity and experience and reinforce societal prejudices. As described above, gender is a complex and dynamic concept that cannot be essentialized to specific identities or roles. As such, we stress that while our analyses identify biological correlates for gender in children, they do not provide evidence for gender essentialism. Our results demonstrate that the association between functional brain organization and gender can be reliably captured, suggesting that complex and nuanced social and environmental factors may jointly influence both brain organization and gender.

**Future Work and Open Questions.** Biomedical research studies to date have largely focused on the influences of sex on brain biology and behavior, without consideration of how gender might contribute to those relationships. Large-scale neuroimaging (and other biomedical) datasets typically lack data pertaining to gender and conflate *sex* and *gender.* As such, there is still a considerable gap in our understanding of the neuroscience of gender, and the intersectional associations between sex, gender, brain, and behavior. Future work in this area, using the ABCD (and other) datasets, should critically investigate open questions addressing this gap. It is important to also recognize that gender is not static and can evolve within an individual over time. As these children get older, their genders are likely to change, along with their brain development. Future analyses should investigate whether individual gender scores and the neurobiological correlates of sex and gender identified here remain consistent throughout adolescence in this cohort.

**A note on *prediction.*** A central goal in neuroscience is to understand how individual differences in the brain are associated with individual differences in sociodemographics (e.g., sex and gender) and behaviors (e.g., cognition and psychopathology)^13^. In machine learning and data science, ‘predictive models' refer to algorithms that *establish associations between independent and dependent variables in existing data, and use those learned associations to “predict” trends and properties in unobserved samples^14^.* In the brain sciences, these models are used to capture associations between brain and behavior in a large sample or dataset and subsequently “predict”, or rather estimate, those behaviors in previously unseen samples based on each individual’s brain data. Our use of the terms ‘prediction’ and ‘predictive model(s/ing)’ in the main text are not intended to suggest that brain functional connectivity patterns at present can ‘predict’ or ‘forecast’ an individual’s sex or gender in the future, or even at present. As stated in our main text, sex and gender are both influenced by biological and social constructs. Our ability to capture meaningful associations between brain functional connectivity and sex and gender suggests that these complex social constructs have reciprocal and compounding influences on brain development and organization.
